# Molecular rhythm alterations in prefrontal cortex and nucleus accumbens associated with opioid use disorder

**DOI:** 10.1101/2021.10.07.463568

**Authors:** Xiangning Xue, Wei Zong, Jill R. Glausier, Sam-Moon Kim, Micah A. Shelton, BaDoi N. Phan, Chaitanya Srinivasan, Andreas R. Pfenning, George C. Tseng, David A. Lewis, Marianne L. Seney, Ryan W. Logan

## Abstract

Severe and persistent disruptions to sleep and circadian rhythms are common features of people with opioid use disorder (OUD). Preclinical findings suggest altered molecular rhythms in the brain are involved in opioid reward and dependence. However, whether molecular rhythms are disrupted in brains of people with OUD remained an open question, critical to understanding the role of circadian rhythms in opioid addiction. We previously used subjects’ times of death (TOD) as a marker of time of day to investigate transcriptional rhythm alterations in psychiatric disorders. Using TOD and RNA sequencing, we discovered rhythmic transcripts in both the dorsolateral prefrontal cortex (DLPFC) and nucleus accumbens (NAc), key brain areas involved in opioid addiction, were largely distinct between OUD and unaffected comparison subjects. Further, fewer rhythmic transcripts were identified in DLPFC of OUD subjects compared to unaffected subjects, but nearly double the number of rhythmic transcripts were found in the NAc of OUD subjects. In OUD, rhythmic transcripts in the NAc peaked either in the evening or near sunrise, and were associated with dopamine, opioid, and GABAergic neurotransmission. Co-expression network analysis identified several OUD-specific modules in the NAc, enriched for transcripts involved in the modulation of dopamine and GABA synapses, including glutamatergic signaling and extracellular matrices. Integrative analyses with human GWAS revealed that rhythmic transcripts in DLPFC and NAc were enriched for genomic loci associated with sleep duration and insomnia. Overall, our results connect transcriptional rhythm changes in dopamine, opioid, and GABAergic synaptic signaling in human brain to sleep-related phenotypes and OUD.

## Introduction

Despite the enormous public health impact of opioids, understanding of the mechanisms which contribute to dependence, abuse, and relapse are lacking. Current pharmacotherapies are largely aimed at symptomatic relief (e.g., mitigating ‘flu-like’ symptoms during acute withdrawal). Remarkably, ~90% of patients with opioid use disorder (OUD) relapse within 12-36 months of beginning treatment^1^. Knowledge of the mechanisms contributing to opioid reinforcement and relapse is critical for designing more effective therapeutics to treat OUD.

Among the most common features associated with OUD are severe and persistent disruptions to sleep and circadian rhythms (e.g., sleep-wake cycles, sleep quality and architecture, corticosterone and melatonin rhythms), which are speculated to foster both craving and relapse^2, 3^. Many individuals (~60%) with OUD have comorbid sleep and circadian disorders and most individuals with a history of opioid use and dependence experience sleep loss, poor sleep quality, and altered sleep-wake cycles^4, 5^. Opioids dose-dependently alter sleep-wake cycles, body temperature and hormonal rhythms^5, 6^. Sleep and circadian disturbances frequently emerge during acute opioid withdrawal and accompany intense cravings and negative affective states^7^. With prolonged opioid use, these sleep and circadian disruptions become more severe^8–10^ and may even intensify cravings and mood disturbances^11, 12^, suggesting that opioids impact sleep and circadian rhythms, which, in turn, contribute to opioid reinforcement and vulnerability to relapse. In fact, craving intensity positively correlates with the severity of sleep and circadian disruptions^13, 14^.

Disruptions to molecular rhythms are also associated with opioid dependence and may be involved in opioid-induced cellular and behavioral plasticity. In nearly every cell in the body, molecular rhythms are controlled by transcriptional—translational feedback loops driven by several transcription factors^15^. The circadian transcription factors CLOCK and BMAL1 dimerize and bind to enhancer elements in promoters to drive rhythmic gene expression. Although estimates vary, ~40-80% of the genome appears to be rhythmically expressed in a tissue and cell-type specific manner^16^, including within brain regions associated with drug reinforcement, craving, and relapse^17, 18^. In individuals with OUD, molecular rhythms in lymphocytes from blood are significantly blunted^19^. Animal studies reveal similar disruptions to molecular rhythms in brain and other tissues following opioid administration^6, 20, 21^. In animals, molecular rhythm disruptions in brain regions associated with OUD also lead to altered locomotor rhythms and increased drug reinforcement and relapse^22^. The entrainment of molecular rhythms in the brain to opioid administration may be involved in drug-seeking and craving^13, 14^. On the other hand, certain variants in canonical circadian genes also predict risk for substance use and substance-associated sleep and circadian disruptions^23, 24^. Together, these findings suggest complex bidirectional relationships between circadian rhythms and opioids—opioids disrupt clocks, which contribute to the development and progression of opioid tolerance and dependence, while disrupted clocks may contribute to the risk for opioid use and dependence.

Human neuroimaging findings suggest that dysfunction of corticostriatal circuits, particularly between the dorsolateral prefrontal cortex (DLPFC) and nucleus accumbens (NAc), are associated with increased risk for substance use^23, 24^. In humans, the DLPFC is involved in cognitive and emotional control, whereas the NAc regulates goal-directed behavior and reward-seeking. In OUD, dysfunction within the DLPFC—NAc circuitry is related to the severity of cognitive impairment and risk for relapse^25^. Emerging evidence also suggests disruptions to molecular rhythms in corticostriatal circuits are involved in opioid-induced cellular and synaptic plasticity. For example, disruption of molecular rhythms in cortical and striatal brain areas promotes tolerance to opioids and enhances opioid withdrawal^26–29^.

To investigate whether molecular rhythms were altered in the brains of subjects with OUD, we assessed molecular rhythms in DLPFC and NAc of unaffected comparison subjects and subjects with OUD using our previous dataset^30^. We found robust transcriptional rhythms in DLPFC and NAc in unaffected subjects that were significantly altered in subjects with OUD. The timing of peak transcript expression differed between unaffected and OUD subjects and brain region. In OUD, alterations in transcriptional rhythms revealed pathways associated with circadian regulation of dopamine, opioid, and GABAergic signaling in DLPFC and NAc. Finally, we discovered genetic associations between transcriptional rhythm changes in DLPFC and NAc in OUD and sleep-related traits.

## Materials and Methods

### Human Subjects

Brains were obtained, following consent from the next-of-kin, during autopsies conducted at the Allegheny County (Pittsburgh, PA; N=39) or the Davidson County (Nashville, TN; N=1) Medical Examiner’s Office. An independent committee of clinicians made consensus, lifetime DSM-IV diagnoses for each subject using the results of a psychological autopsy, including structured interviews with family members and review of medical records, and toxicological and neuropathological reports^31^. The same approach was used to confirm the absence of lifetime psychiatric and neurologic disorders in the unaffected comparison subjects. All procedures were approved by the University of Pittsburgh Committee for Oversight of Research and Clinical Training Involving Decedents and Institutional Review Board for Biomedical Research. Each OUD subject (n=20) was matched with an unaffected comparison subject (n=20) for sex and as closely as possible for age^30^ (**Table S1-S2)**. Cohorts differed by race composition (p=0.02) and brain pH (p=0.015; mean difference was 0.2 pH units), but did not differ in PMI, age, RIN, pH, or TOD (p>0.25). DLPFC area 9 and NAc were identified and collected as previously described^30^.

### Rhythmicity analyses

TOD of each subject was adjusted to internal biological clock time by normalization to zeitgeber time (ZT). Cosinor fitting was used to detect rhythmicity of transcripts expression. Sinusoidal curves were fitted using the nonlinear least-squares method, with coefficient of determination (*R*^2^) used as proxy of goodness-of-fit. An estimate of the empirical p-value was determined using a null distribution of *R*^2^ generated from 1000 TOD-randomized expression datasets. Molecular rhythms were first assessed separately in unaffected comparison subjects and subjects with OUD. Rhythmic transcripts were compared using significance cutoffs (p<0.05) and a threshold-free approach (rank-rank hypergeometric overlap (RRHO))^32^. Transcripts with OUD-related differences in rhythmicity were determined using the difference in *R*^2^. Transcripts with *ΔR*^2^>0 when 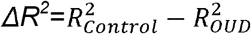 were defined as being significantly less rhythmic in OUD. Transcripts with *ΔR*^2^>0 when 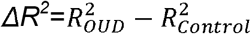 were defined as being significantly more rhythmic in OUD. We generated a null distribution of *ΔR^2^* using the null *R^2^* values permuted in unaffected comparisons and OUD subjects separately. Any transcript with significant *ΔR*^2^ (p<0.05 through permutation test) are denoted as having a significantly less or more rhythmicity in OUD. We further restrict the change in rhythmicity analysis to transcripts that are significantly rhythmic in one group or the other. For a transcript to be considered less rhythmic in OUD, it had to: 1) be significantly rhythmic in unaffected comparisons (p<0.05); 2) be significantly less rhythmic in OUD (p<0.05). For a transcript to be more rhythmic in OUD, it had to: 1) be significantly rhythmic in OUD (p<0.05); 2) be significantly more rhythmic in OUD (p<0.05). We also assessed differences in phase, amplitude, or base; analyses were restricted to transcripts significantly rhythmic in both unaffected comparisons and OUD subjects.

### Heatmaps

Transcript expression levels were Z-transformed and ordered by their phase value (peak hour). Each column represents a subject, ordered by TOD. We generated heatmaps for 1) top 200 rhythmic transcripts in unaffected comparison subjects; 2) top 200 rhythmic transcripts identified in unaffected comparison subjects but plotted for OUD subjects; 3) top 200 rhythmic transcripts identified in OUD subjects; 4) top 200 rhythmic transcripts identified in OUD subjects, but plotted for unaffected comparison subjects.

### Scatter plots

Scatter plots were generated to represent transcript expression rhythms. TOD on ZT scale is indicated on the x axis and transcript expression level on the y axis, with each dot indicating a subject. The red line is the fitted sinusoidal curve. For each brain region, scatter plots were generated for the top three transcripts that were significantly less rhythmic in OUD subjects and the top three transcripts that were significantly more rhythmic in subjects with OUD relative to unaffected comparison subjects.

### RNA Sequencing analyses

Overrepresentation of pathways (GO, KEGG, Hallmark, Canonical Pathways, Reactome, BioCarta, CORUM) was assessed using Metascape (http://www.metascape.org) for rhythmic transcripts in DLPFC and NAc, transcripts that were more rhythmic or less rhythmic in OUD, and transcripts in co-expression networks, with expressed transcripts as background. Networks were visualized with Cytoscape. INGENUITY^®^ Pathway Analysis (Qiagen) was used to predict upstream regulators of rhythmic transcripts. Rank-rank hypergeometric overlap (RRHO)^32, 34^ was used to assess overlap of rhythmic transcripts between unaffected comparison subjects and subjects with OUD.

### Identification of OUD-specific co-expression networks

We used weighted gene co-expression network analysis (WGCNA) to identify transcript modules across samples^35^. Networks were built separately in each brain region and disease group. We used Fisher’s exact test to determine enrichment of rhythmic transcripts or transcripts that were significantly less rhythmic or more rhythmic in OUD subjects within each of the WGCNA modules. ARACNe was used to identify hub and OUD-specific hub transcripts for network analysis^36^ and Cytoscape was used to visualize networks. Overrepresentation of pathway categories for each module was assessed using Metascape, with the 5000 WGNCA-analyzed transcripts as background.

### Integration of rhythmic transcripts with GWAS

Transcripts that were rhythmic within disease groups as well as transcripts that were less rhythmic or more rhythmic in OUD subjects (corrected p<0.05) were used to construct foregrounds for GWAS enrichment. We computed the partitioned heritability (GWAS enrichment) of noncoding regions containing and surrounding OUD rhythmic transcript sets using the LD score regression pipeline for enrichment^37^. LD score regression coefficients were adjusted for FDR<0.05 on enrichments performed on each of the included GWAS foregrounds. A significant p-value indicates enrichment of the foreground genomic regions for GWAS single nucleotide polymorphisms relative to the background.

## Results

### Distinct transcriptional rhythms in subjects with OUD

Given the convergent evidence in clinical and preclinical evidence of altered circadian rhythms in opioid addiction^7, 38, 39^, we investigated whether brain transcriptional rhythms were also altered in DLPFC and NAc, brain regions strongly implicated in OUD^40–43^. In DLPFC, we identified fewer rhythmic transcripts in subjects with OUD (n = 339) compared to unaffected comparison subjects (n = 730) (**Table S3** and **Table S4**)), with only 19 rhythmic transcripts shared between groups (**Fig.1A**). Results using a threshold-free approach, RRHO, further supported a lack of overlap between unaffected and OUD subjects in rhythmic transcripts in DLPFC (**Fig.1B**). The top 250 rhythmic transcripts peaked across the day in unaffected comparison subjects, but these same transcripts were arrhythmic in subjects with OUD (**Fig.1C**). Similarly, the top rhythmic transcripts in the DLPFC of OUD subjects were arrhythmic in unaffected comparison subjects (**Fig.1D**).

**Figure 1.**
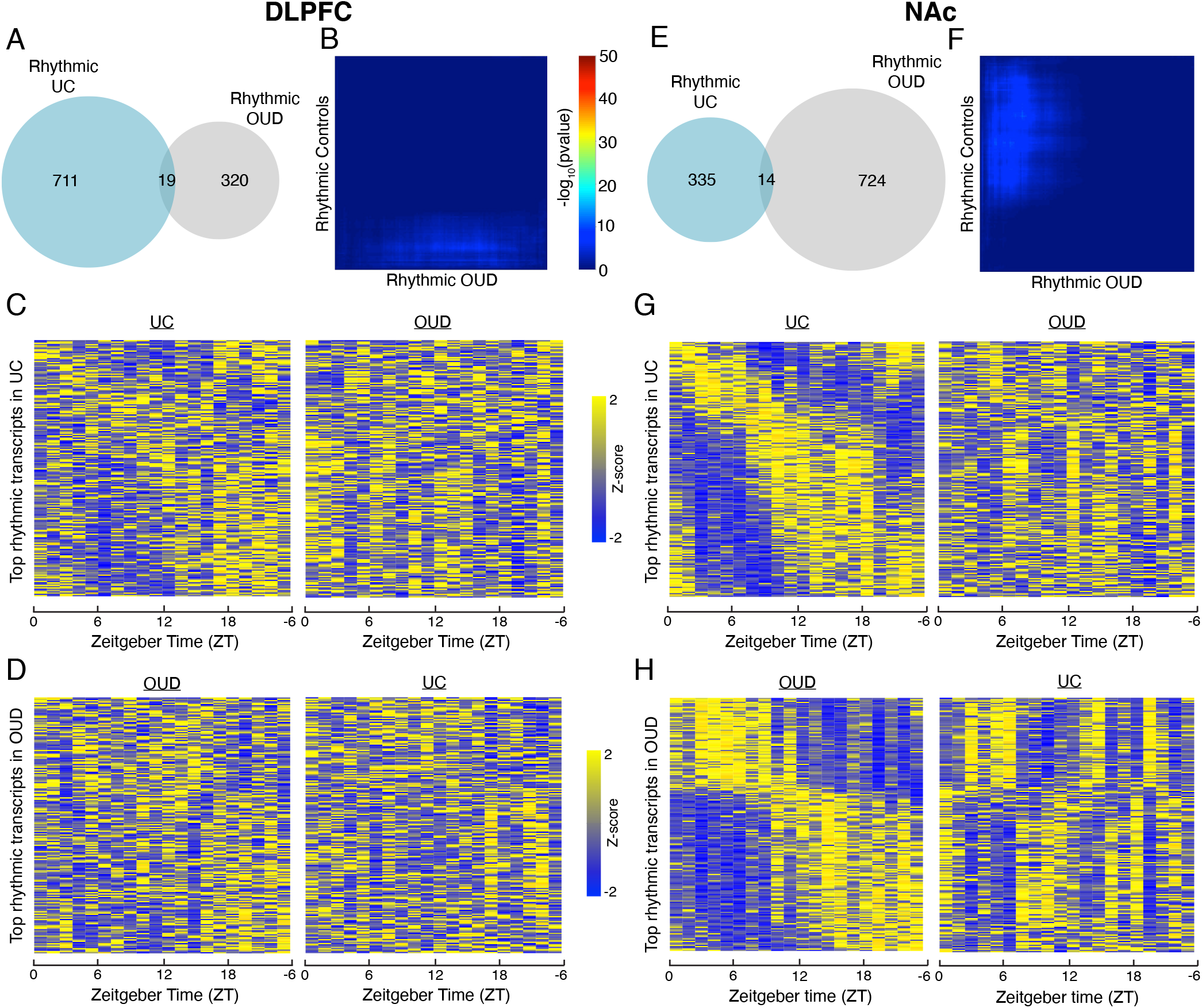
Rhythmic transcripts are largely distinct in unaffected comparison subjects and subjects with opioid use disorder. **A.** In the dorsolateral prefrontal cortex (DLPFC), there were 730 rhythmic transcripts detected in unaffected comparison (UC) subjects and 339 in subjects with opioid use disorder (OUD). Notably, only 19 transcripts were rhythmic in both UC subjects and subjects with OUD. **B.** Rank-rank hypergeometric overlap was used as a threshold-free approach to confirm the lack of overlap in rhythmicity patterns in the DLPFC of UC subjects and subjects with OUD. **C.** Heatmap of the top 200 circadian transcripts identified in the DLPFC of UC subjects (left), with transcripts peaking across the day. Expression levels are Z-transformed for each transcript, and the transcripts are ordered by their circadian phase value (peak hour). Each column represents a subject and the subjects are ordered by time of death. The top 200 rhythmic transcripts identified in UC subjects are then plotted for subjects with OUD (right), indicating disrupted rhythmicity of normally rhythmic transcripts in subjects with OUD. **D.** The top 200 rhythmic transcripts identified in OUD subjects in the DLPFC (left) are then plotted in UC subjects (right). **E.** In the nucleus accumbens (NAc), there were 349 rhythmic transcripts detected in UC subjects and 738 in subjects with OUD. Notably, only 14 transcripts were rhythmic in both UC subjects and subjects with OUD. **F.** Rank-rank hypergeometric overlap was used as a threshold-free approach to confirm the lack of overlap in rhythmicity patterns in the NAc of UC subjects and subjects with OUD. **G.** Heatmap for the top 200 circadian transcripts identified in the NAc of UC subjects (left). The top 200 rhythmic transcripts identified in UC subjects are then plotted for subjects with OUD (right). **H.** The top 200 rhythmic transcripts identified in OUD subjects in the NAc (left) are then plotted in UC subjects (right).

We performed a similar analysis of transcriptional rhythmicity in the NAc of unaffected comparison and OUD subjects. Surprisingly, OUD subjects had more than twice as many rhythmic transcripts compared to unaffected comparison subjects (738 and 349, respectively) (**Table S5** and **Table S6**), with an overlap of only 14 transcripts (**Fig.1E**). RRHO further supported an overall lack of overlap between unaffected comparison and OUD subjects in the NAc (**Fig.1F**). In unaffected comparison subjects, the top 250 rhythmic transcripts peaked across the day, while these transcripts were arrhythmic in subjects with OUD (**Fig.1G**). In OUD subjects, the following patterns of diurnal expression were identified in top rhythmic transcripts: 1) peaks of expression during the day and troughs at night; or 2) troughs of expression during the day and peaks at night (**Fig.1H**). Using the top 250 rhythmic transcripts in NAc of OUD subjects, these anti-phasic patterns of transcriptional rhythms were present in subjects with OUD and appear to exhibit near 12-hour rhythms of expression in unaffected comparison subjects compared to the 24-hour rhythm in subjects with OUD (**Fig.1H**).

### Distinct peak times of rhythmic transcripts in subjects with OUD

Having identified rhythmic transcripts in the DLPFC and NAc of unaffected comparison and OUD subjects, we next evaluated the timings of peak transcript expression. In DLPFC, we observed two peaks of expression in unaffected comparison subjects at ~ZT4 and another at ~ZT16 (**Fig.2A, left**), nearly 12 hours apart from each other; in other words, transcripts tended to peak at either ZT4 or ZT16. In contrast to unaffected comparison subjects, rhythmic transcripts in the DLPFC of OUD subjects did not exhibit distinct peaks of expression (**Fig.2A, right**). In NAc of unaffected comparison subjects, most of the rhythmic transcripts peaked ~ZT10 (**Fig.2B, left**). Rhythmic transcripts in the NAc of OUD subjects peaked at either ~ZT11 and ~ZT23 (**Fig.2B, right**).

**Figure 2.**
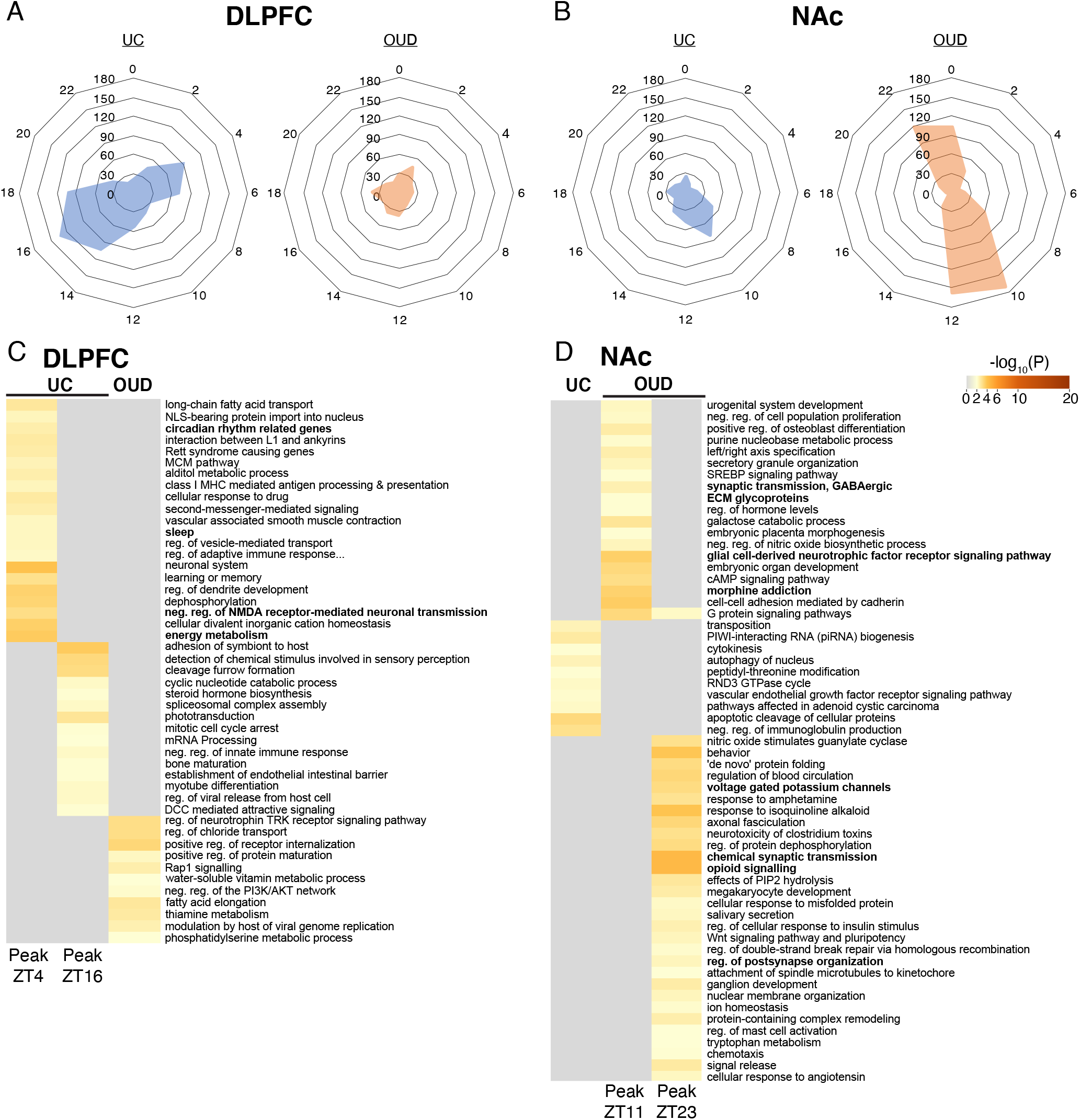
Distinct peak times for rhythmic transcripts in subjects with opioid use disorder (OUD) compared to unaffected comparison (UC) subjects. **A.** In the dorsolateral prefrontal cortex of UC subjects, transcripts generally peaked at either ZT4 or ZT16, approximately 12 hours apart. Rhythmic transcripts in the DLPFC in subjects with OUD did not peak at consistent times. **B.** In the nucleus accumbens (NAc), rhythmic transcripts in UC subjects generally peaked at ZT10. Rhythmic transcripts in the NAc of subjects with OUD peaked at either ZT11 or ZT23, approximately 12 hours apart. **C.** Transcripts peaking at ZT4 in the DLPFC of UC subjects were enriched for pathways related to rhythms (e.g., circadian rhythm related genes, sleep) and neurotransmission (e.g., negative regulation of NMDA receptor-mediated neuronal transmission), while transcripts peaking at ZT12 were enriched for immune-related pathways (e.g., adhesion of symbiont to host, negative regulation of innate immune response). Rhythmic transcripts in the DLPFC of OUD subjects were enriched for regulation of neurotrophin TRK receptor signaling pathway and positive regulation of receptor internalization. **D.** Rhythmic transcripts in the NAc of OUD subjects were enriched for apoptotic cleavage of cellular proteins. In the NAc of OUD subjects, rhythmic transcripts peaking at ZT11 were enriched for morphine addiction, glial cell-derived neurotrophic factor receptor signaling, ECM glycoproteins, and synaptic transmission, GABAergic, while transcripts peaking at ZT23 were enriched for opioid signaling, voltage gated potassium channels, and synapse-related pathways (e.g., regulation of postsynapse organization, chemical synaptic transmission).

The observation that rhythmic transcripts form two separate clusters with peaks ~12-hours apart in DLPFC of comparison subjects and in NAc of OUD subjects provided justification for further investigation of the biological pathways represented by these clusters of transcripts. In DLPFC of unaffected comparison subjects, transcripts that peak at ZT4 (approximately midmorning to noon) were enriched for regulation of ion transport, energy metabolism, and negative regulation of NMDA receptor-mediated neuronal transmission, and circadian rhythm-related genes (**Fig.2C**). Transcripts that peak at ZT16 (approximately late evening to midnight) in the DLPFC of unaffected comparison subjects were enriched for immune-related pathways (*e.g*., adhesion of symbiont to host, negative regulation of viral transcription; **Fig.2C**). Rhythmic transcripts in the DLPFC of OUD subjects were enriched for pathways related to receptor internalization and neurotrophin signaling (**Fig.2C**). In the NAc of unaffected comparison subjects, rhythmic transcripts were enriched for immune pathways (*e.g*., negative regulation of immunoglobulin production) and small noncoding RNAs (*e.g*., PIWI-interacting (piRNA) RNA biogenesis). Additionally, we investigated pathways associated with rhythmic transcripts peaking at either ~ZT11 or ~ZT23 in NAc of OUD subjects. Interestingly, transcripts peaking at ZT11 and ZT23 were both enriched for opioid-related signaling pathways (**Fig.2D**). Transcripts peaking at ~ZT11 (evening) in the NAc of OUD subjects were enriched for morphine addiction, synaptic transmission, GABAergic transmission, and glial cell-derived neurotrophic factor receptor signaling pathways. The transcripts that peaked at ~ZT23 (right before sunrise) were enriched for chemical synaptic transmission, voltage gated potassium channels, and opioid signaling (**Fig.2D**). Collectively, these findings indicate rhythmic transcripts: 1) in the DLPFC of unaffected comparison subjects were primarily associated with immune and excitatory synaptic signaling; and 2) in the NAc of OUD subjects were associated with opioid signaling and GABAergic neurotransmission.

### Alterations in transcriptional rhythmicity in OUD

Given that we observed very little overlap of transcripts that were rhythmic in both unaffected comparison and OUD subjects, we decided to empirically test whether any transcripts were significantly less rhythmic or more rhythmic in OUD subjects, different from assessing overlap of rhythmic transcripts using arbitrary statistical cutoffs. In DLPFC, we identified 548 transcripts that were significantly less rhythmic in subjects with OUD relative to unaffected comparison subjects (**Table S7**). Several of top rhythmic transcripts in DLPFC that were less rhythmic in OUD subjects included *APBA2* (amyloid beta precursor protein binding family A member 2)*, FAT3* (FAT atypical cadherin 3), and *AC083798.2* (Long non-coding RNA, LncRNA) (**Fig. 3A**). The top biological pathways associated with transcripts that were less rhythmic in OUD were Netrin and Eicosanoid signaling (**Fig.3C**) and the top predicted upstream regulators included the canonical circadian proteins, PER1 and PER2 (**Fig.3C**), suggesting molecular clock disruptions in DLPFC of OUD subjects.

**Figure 3.**
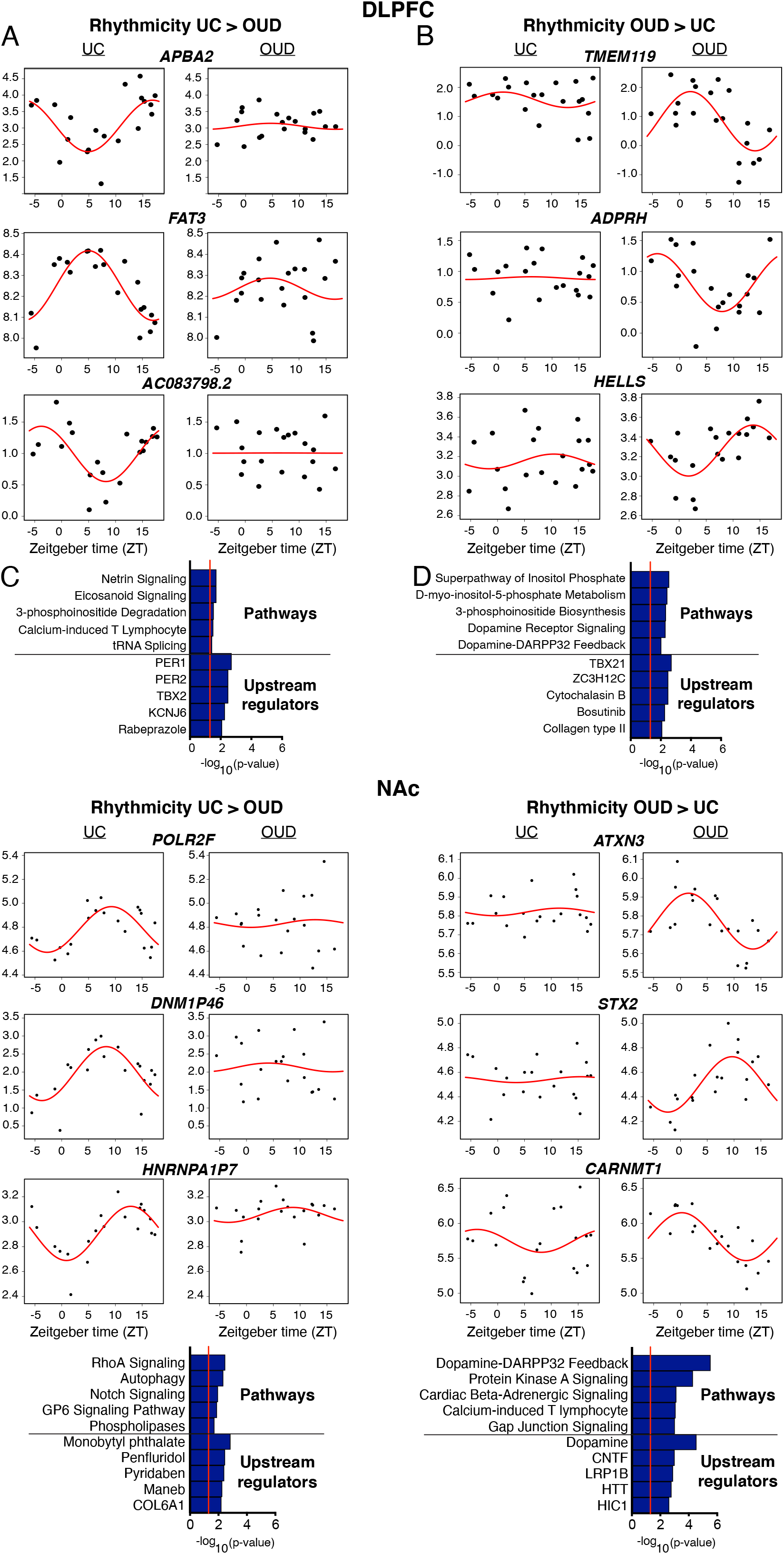
Scatterplots indicating rhythmicity for transcripts that were significantly more or less rhythmic in the dorsolateral prefrontal cortex (DLPFC) in subjects with opioid use disorder (OUD) compared to unaffected comparison subjects. Each dot indicates a subject with x axis indicating the time of death (TOD) on ZT scale (−6~18h) and y axis indicating transcript expression level. The red line is fitted sinusoidal curve. **A.** Scatterplots indicating rhythmicity of *APBA2, FAT3*, and *AC083798.2* in unaffected comparison subjects (left), which are significantly less rhythmic in subjects with OUD (right). **B.** The top pathways represented by transcripts that are less rhythmic in the DLPFC in OUD are related to netrin signaling and eicosanoid signaling, and the top predicted upstream regulators are PER1 and PER2. **C.** Scatterplots indicating lack of rhythmicity of *TMEM119, ADPRH*, and *HELLS* in unaffected comparison subjects (left), and these transcripts are significantly more rhythmic in subjects with OUD (right). **D.** The top pathways represented by transcripts that are more rhythmic in the DLPFC in OUD are related to inositol and dopamine, and the top predicted upstream regulators are TBX21 and ZC3H12C.

We also identified 209 transcripts that were significantly more rhythmic in DLPFC of subjects with OUD compared to unaffected comparison subjects, with *TMEM119, ADPRH*, and *HELLS* as the top three transcripts (**Fig.3B; Table S8**). The top pathways associated with transcripts that were more rhythmic in OUD subjects in the DLPFC included inositol biosynthesis and metabolism, along with dopamine receptor signaling and DARPP32 feedback (**Fig.3D**), with TBX21 and ZC3H12C as top predicted upstream regulators.

In NAc, we found 406 transcripts that were significantly more rhythmic in unaffected comparison subjects compared to subjects with OUD, including *POLR2F, DNM1P46*, and *HNRNPA1P7* (**Fig.3E**; **Table S9**). From these transcripts, we also identified several interacting signaling pathways including RhoA^44^, Notch^45^, and GP6^46^ (**Fig.3G**). Upstream regulators included several pesticide agents with neurotoxic profiles (*e.g*., monobutyl phthalate) and penfluridol, an antipsychotic medication with primary action at the dopamine 2 receptor^47^. Among the 762 transcripts that were significantly more rhythmic in the NAc of subjects with OUD compared to unaffected comparison subjects, the top transcripts included *ATXN3, STX2*, and *CARNMT1* (**Fig.3F**; **Table S10**). Like DLPFC, we identified dopamine-DARPP32 feedback pathways among the top enriched pathways along with dopamine as the top predicted upstream regulator of transcripts that were more rhythmic in NAc of OUD subjects (**Fig.3H**). Other top pathways included protein kinase A, beta-adrenergic, and gap junction signaling, along with the calcium-induced T lymphocyte pathway (**Fig.3H**). Overall, our findings suggest pathways related to dopamine signaling were more rhythmic in both DLPFC and NAc of OUD subjects.

### Gene module enrichment of synapse-related and glycoprotein signaling in OUD

WGCNA identified 16 modules in DLPFC of unaffected subjects with only one module enriched for rhythmic transcripts and only one module enriched for transcripts that were significantly less rhythmic in OUD (**Fig.S1A**). Additionally, we identified 20 modules in DLPFC of OUD subjects and none of these modules were enriched for rhythmic transcripts or for transcripts that were significantly more or less rhythmic in OUD.

From the 19 modules identified in the NAc of unaffected subjects, only one module was enriched for transcripts that were less rhythmic in the NAc of OUD subjects (**Fig.S1B**). In OUD subjects, we identified 16 modules in the NAc and three of these modules (brown, red, pink) were enriched for rhythmic transcripts. Both red and pink modules were enriched for transcripts that were more rhythmic in the NAc of OUD subjects (**Fig.4A**). Several transcripts that are key regulators of synaptic signaling were present in each of the modules that were significantly enriched for transcripts that were more rhythmic in the NAc of OUD subjects. These transcripts included: *GRIN3A*^48, 49^*, SLC6A7^50^, KCNJ6*^51, 52^, *GABRQ*^53^, and *HPCAL1^54^* (brown module); *SEMA5B*^55^ and *SHISA6*^56^ (pink module); and *PCP4^57^* and *PPP1R1B (DARPP-32)^58^* (red module). Pathway enrichment analyses of transcripts comprising the networks in brown, pink, and red modules further supported the connection between rhythmic transcripts in OUD and synaptic function in the NAc, including neurotransmitter receptors and postsynaptic signal transmission, trans-synaptic signaling, positive regulation of excitatory postsynaptic potential, and pathways related to extracellular matrices (ECM) and brain morphology (*e.g*., ECM glycoproteins, cell-cell adhesion molecules, and axon development) (**Fig.4B**).

**Figure 4.**
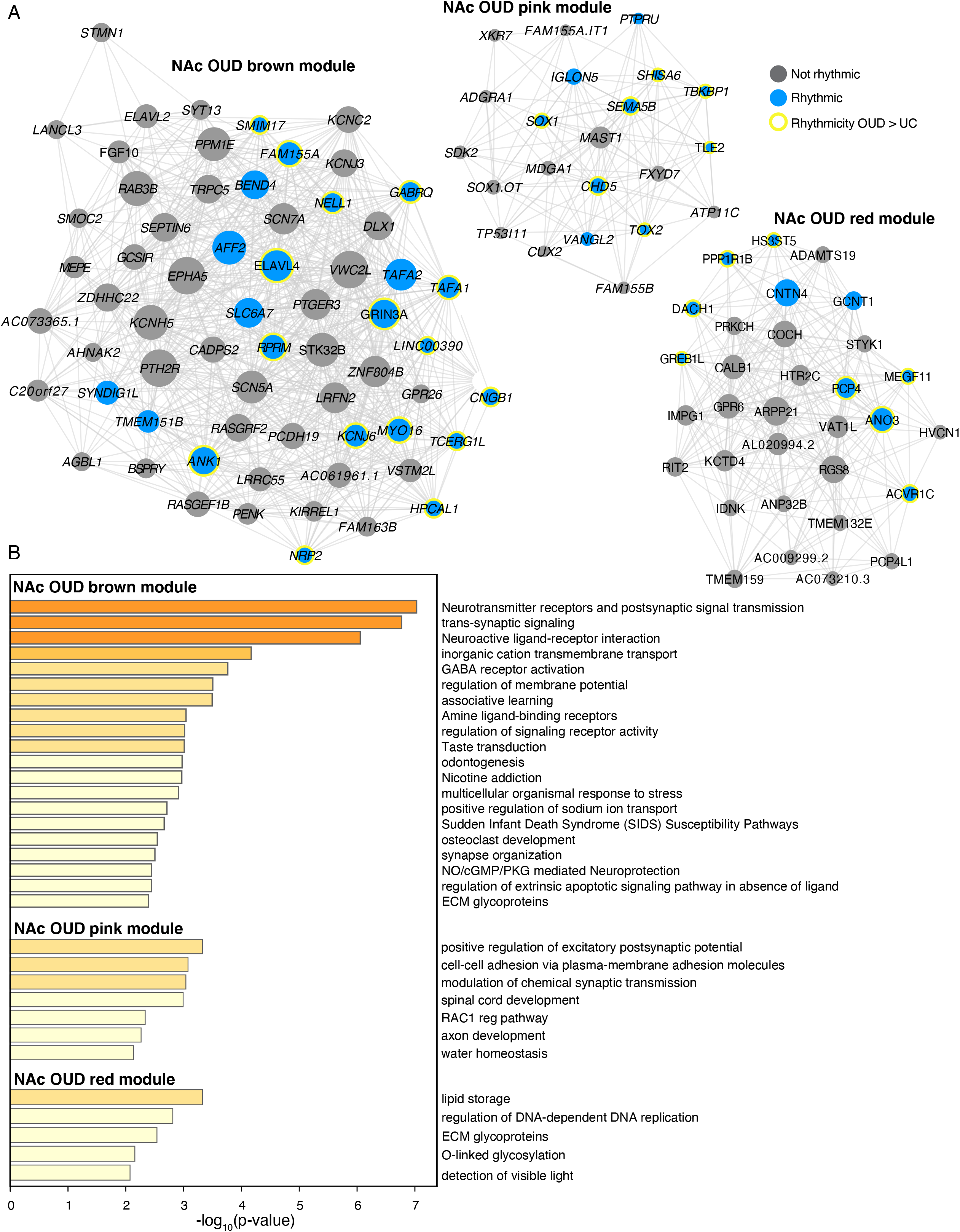
OUD associated gene networks in the NAc. **A.** Weighted gene co-expression network analysis (WGCNA) was used to generate co-expression modules, with the network structure generated on each brain region separately. The identified modules that survived module preservation analysis were arbitrarily assigned colors and module differential connectivity (MDC) analysis compared the identified modules in OUD and unaffected comparison subjects. **A.** MDC analysis indicated a gain of connectivity in the NAc for the brown, pink, and red modules. Node size indicates the degree of connectivity for that transcript. Blue nodes indicate rhythmic transcripts and yellow halos indicate transcripts that were significantly more rhythmic in OUD. Edges indicate significant co-expression between two particular transcripts. **B.** Pathway enrichment analysis within the NAc OUD brown module, the NAc OUD pink module, and the NAc OUD red module. Warmer colors indicate increasing -log_10_ p-value.

### Brain region-specific genetic associations between altered transcriptional rhythms and sleep phenotypes

Given our findings that reveal altered transcriptional rhythmicity in the brains of subjects with OUD, we investigated whether transcripts with significant alterations in rhythmicity in the DLPFC or NAc of OUD subjects were associated with GWAS identified genetic variants of sleep-related traits. To test this idea, we used GWAS studies to integrate significant genomic loci from various sleep traits (*e.g*., insomnia, morningness, and sleep duration). Genomic loci identified by GWAS overlap with intronic and distal intergenic non-coding regions within *cis*-acting regulators of gene expression^59^. Using the intronic and distal intergenic regions, we examined whether these genomic regions proximal to our rhythmic transcripts in unaffected comparison subjects and subjects with OUD were enriched for genetic associations with sleep-related traits. We identified significant enrichments for rhythmic transcripts in DLPFC of unaffected subjects for insomnia and long sleep duration (**Fig.5A**). Transcripts that were more rhythmic in the DLPFC of unaffected subjects compared to OUD subjects were significantly enriched in insomnia and morningness GWAS (**Fig. 5A**). In NAc of OUD subjects, highly rhythmic transcripts were significantly enriched for total sleep duration GWAS, including transcripts that were more rhythmic in OUD compared to unaffected subjects (**Fig.5B**). Together, our integrative analyses of rhythmic transcriptomes with human GWAS establishes connections between alterations of transcriptional rhythms in corticostriatal circuitry, opioid addiction, and phenotypes associated with sleep disturbances (*i.e*., shorter sleep durations and insomnia).

**Figure 5.**
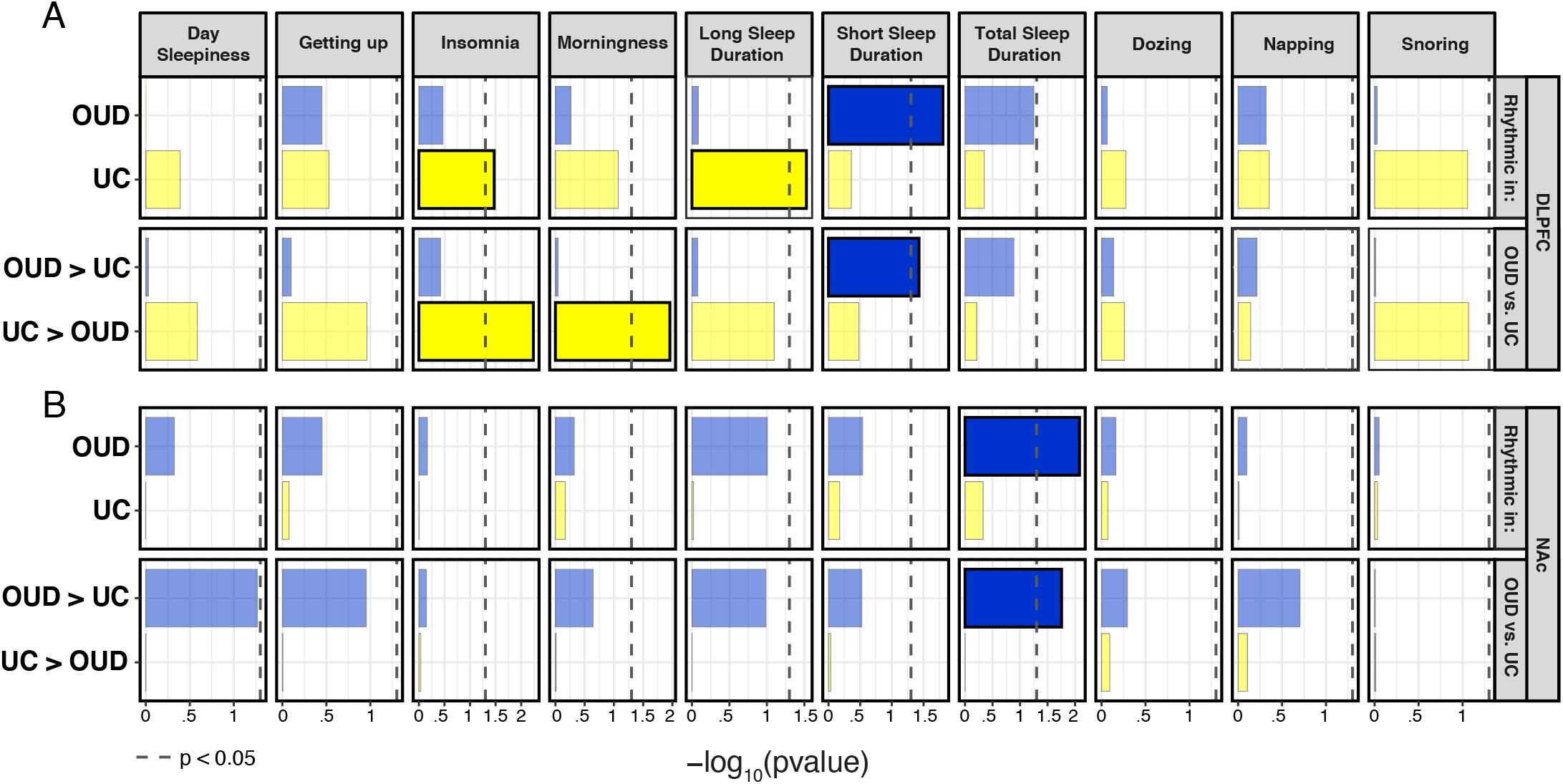
Rhythmic transcripts in the dorsolateral prefrontal cortex (DLPFC) and nucleus accumbens (NAc) enrich for genetic associations with sleep-related traits. Genome-wide associated studies (GWAS) have identified loci associated with various sleep-related traits. We investigated whether rhythmic transcripts as well as transcripts that were significantly more or less rhythmic in subjects with opioid use disorder (OUD) were enriched for genetic associations with sleep-related traits. **A.** In the DLPFC of unaffected comparison (UC) subjects, there was significant enrichment of rhythmic transcripts for genes associated with insomnia and long sleep duration. Insomnia and morningness were associated with transcripts that were significantly less rhythmic in DLPFC of OUD subjects. In the DLPFC of OUD subjects, rhythmic transcripts and transcripts that were more rhythmic in OUD subjects in DLPFC were enriched for genes associated with short sleep duration. **B.** In NAc of OUD subjects, there was enrichment of rhythmic transcripts and transcripts that were more rhythmic in OUD in total sleep duration. There were no significant associations in the NAc of UC subjects.

## Discussion

Our data demonstrate robust transcriptional rhythms in both the DLPFC and NAc of unaffected subjects and subjects with OUD. Rhythmic transcripts in unaffected subjects were largely distinct from those in subjects with OUD, suggesting that chronic opioid use leads to the emergence of rhythmicity in specific transcripts involved in corticostriatal circuit function. In line with this idea, many of the transcripts that were rhythmic in the DLPFC and NAc of OUD subjects were enriched for pathways related to the regulation of dopamine neurotransmission. Using integrative analyses that combine diurnal patterns of transcriptional regulation and relevant GWAS, our findings revealed novel gene-trait relationships between transcripts that were significantly more rhythmic in DLPFC and NAc of OUD subjects and sleep phenotypes. Additionally, we identified transcripts that were significantly less rhythmic in OUD subjects, also with significant associations to sleep related GWAS.

Similar to previous studies in human postmortem brain^30, 60^, we found robust transcriptional rhythms in unaffected subjects in both the DLPFC and NAc. In the DLPFC of unaffected subjects, many of the pathways that were enriched from rhythmic transcripts were related to circadian rhythms, sleep, metabolism, immune response, and synaptic and neural transmission. Pathways related to piRNAs, autophagy, and GTPase cycle were among those enriched in the NAc of unaffected subjects. Notably, there was minimal overlap between the transcripts we identified as rhythmic in either the DLPFC or NAc from unaffected subjects compared to OUD subjects. Several pathways which have been previously linked to the effects of opioids were enriched in rhythmic transcripts in the DLPFC of OUD subjects, including neurotrophin TRK receptor signaling^61, 62^ and Rap1 signaling^63^. For example, neurotrophin activation of TRK receptors signals through various molecular cascades (*e.g*., cAMP and ERK) is involved in opioid-induced neural and synaptic plasticity^64, 65^. Dysfunction in neurotrophin TRK receptor signaling and opioid receptor signaling has been associated with psychiatric disorders^62^. Moreover, Rap1 may be involved in a subfamily of GTPase-activating proteins that influence mu-opioid receptor activation^66^ and neurotransmission^63^. Interestingly, Rap1-dependent signaling has been shown to modulate neuronal excitability and drug reward-related behaviors in mice^67^.

In the NAc of OUD subjects, enriched pathways from rhythmic transcripts were related to GABAergic neurotransmission, morphine, opioid signaling, postsynaptic organization, and glial cell neurotrophic factors, along with extracellular matrix (ECM) glycoproteins, among others. Interestingly, we recently reported that transcripts associated with microglial and ECM pathways were differentially expressed in the DLPFC and NAc of subjects with OUD when time of death was not taken into consideration^30^. The current results suggest that these transcripts and their related pathways may be altered at specific times of day. While the functional impact of rhythmic alterations in glia^68^ and brain scaffolding^69, 70^ needs to be explored further in OUD, microglial regulation of neuroinflammation, in addition to the consequences on the ECM and the functional impacts on synaptic physiology may be critically involved in the long-term effects of opioids on the brain^30, 71, 72^.

Many of the rhythmic transcripts we identified in the DLPFC of unaffected subjects and in the NAc of OUD subjects generally peaked at different times of day. In unaffected subjects, rhythmic transcripts tended to peak at either ZT4 (*i.e*., mid-morning) or ZT16 (*i.e*., late evening), represented by distinct sets of enriched pathways. For example, circadian rhythm and sleep related transcripts peak at ZT4, while other pathways peak at ZT16. In OUD subjects, rhythmic transcripts did not exhibit these two peaks, possibly due to this group having fewer than half the rhythmic transcripts compared to the unaffected group. In contrast, rhythmic transcripts in the NAc exhibited two peaks at ZT11 (*i.e*., evening) or ZT23 (*i.e*., prior to “sunrise”) in OUD subjects. Several of the pathways that peaked at ZT11 were related to glia, ECM, morphine signaling, and GABAergic signaling, while the pathways that peaked at ZT23 were potassium channels, synaptic transmission, opioid and Wnt signaling, and others. We also observed hints that transcripts exhibiting a 24-hour rhythm in the NAc of OUD subjects might exhibit ultradian rhythms (*i.e*., less than 24-hours) in unaffected comparison subjects, but we were not powered to determine if these transcripts did indeed exhibit 12-hour rhythms. 12-hour rhythms in neuronal and synaptic activity, neurotransmission (*e.g*., dopaminergic^73^), and behaviors^27–29^ have been described in rodent models^74^.

In subjects with OUD, we found transcripts that were significantly less rhythmic in DLPFC and NAc when compared to unaffected comparison subjects; these transcripts were related to many pathways of brain function, including synapse and immune signaling. For example, the transcript *APBA2* was less rhythmic in DLPFC of OUD subjects and encodes for a synaptic adaptor protein, which when disrupted leads to impaired synaptic formation and vesicle trafficking in excitatory synapses^75^. In addition, *APBA2* variants were associated with impulsivity and addiction vulnerability^76^. APBA2 directly binds neurexin proteins that are neuron-specific surface proteins involved in synaptic formation and netrin signaling^77^. Netrin signaling was the top pathway enriched from transcripts that were less rhythmic in DLPFC of OUD subjects. Other pathways included eicosanoid signaling, involved in synaptic plasticity and inflammation^78^; 3-phosphoinositide degradation, involved in neuronal hyper-excitability and associated with various psychiatric disorders^79^; calcium-induced T lymphocyte, which tunes T-cells to coordinate immune responses^80^; and tRNA splicing^81^. In the NAc of OUD subjects, transcripts that were less rhythmic were related to synapses and substance use. For example, one transcript, *HNRNPA1P7*, belongs to a family of RNA-binding proteins involved in cytoskeletal organization and synaptic activity, and more recently, substance use^82^. Both RhoA and Notch signaling pathways were also enriched for transcripts that were less rhythmic in NAc of OUD subjects, both of which are involved in opioid tolerance and withdrawal^44^.

We also identified many transcripts that were significantly more rhythmic in OUD subjects. For example, *TMEM119*, a robust marker for microglia^83^, was among the top transcripts that were highly rhythmic in the DLPFC of OUD subjects, resembling increased glial reactivity at certain times of day^84^ in OUD. In support of this, *Tmem119* has a robust expression rhythm in the mouse suprachiasmatic nucleus associating with circadian-dependent modulation of glial activity^85^. Many of the top pathways that were enriched among transcripts that were more rhythmic in DLPFC of OUD subjects were related to inositol phosphates, key regulators of cell signaling^86^. The inositol phosphate pathway has been shown to impact cellular and molecular rhythms^87^, suggesting molecular changes in diurnal patterns of expression in the DLPFC of OUD subjects are, in part, driven by alterations in inositol signaling. Other pathways included feedback regulation of dopamine neurotransmission, and the top upstream regulators were TBX21 and ZC3H12C. Both transcription factors modulate dopamine’s actions on immune cells in the brain, controlling the activation of T-cells, consequently regulating the neuroinflammatory response^88^.

In contrast to DLPFC, most transcripts that were significantly more rhythmic in the NAc of OUD subjects were related to synapses. For instance, *ATXN3*, among the top rhythmic transcripts, is involved in the formation of dendritic spines and new synapses in rodent brain^89^ and *STX2*, also among the top rhythmic, regulates vesicle release of neurotransmitters^90, 91^. In addition, *GRIN3A* was significantly more rhythmic in NAc of OUD subjects, and notably, was identified as a hub transcript in the OUD-specific brown module. The brown module was specific to OUD and the NAc, mainly comprised of transcripts involved in neurotransmission, such as postsynaptic receptors, trans-synaptic signaling, neuroactive ligand-receptor signaling, and GABA receptor activation. *GRIN3A* has been to be involved in opioid-induced synaptic plasticity of both excitatory and inhibitory circuits in the NAc following chronic administration^49, 92, 93^ and variants of *GRIN3A* were identified as alleles associated with therapeutic response to methadone in people with opioid addiction^94^. Previously, we found that neuroinflammatory pathways were enriched in differentially expressed transcripts in the NAc from OUD subjects, including interferon (IFN) signaling^30^. Intriguingly, IFN signaling interacts with GRIN3A, whereby elevated IFN induces NMDA-evoked glutamate release^48^. Thus, cycles of opioid withdrawal may elevate IFN levels, leading to changes in excitatory signaling in the NAc and other brain regions, involved in opioid-induced synaptic plasticity and behavioral consequences. Based on our findings in human brains, rhythmicity of GRIN3A-dependent signaling may also be involved in opioid-induced excitatory synaptic plasticity^95^.

We integrated the transcriptional rhythm profiles in human brain with sleep-related GWAS findings to begin to identify novel gene-trait relationships in OUD. Using integrative GWAS analyses, we found that transcripts that were less rhythmic in the DLPFC of OUD subjects were significantly enriched for genomic loci associated with insomnia and morning preference GWAS. Further, transcripts that were more rhythmic in DLPFC of OUD subjects was significantly associated with short sleep duration GWAS. Increased transcriptional rhythmicity in the NAc of OUD subjects were significantly related to total sleep duration. Our findings support associations between genetic risk for sleep alterations, brain-region specific changes in transcriptional rhythmicity, and OUD. Given the roles for both of the DLPFC^96^ and NAc^97–99^ in sleep and substance use, our results provide novel putative mechanisms of transcriptional rhythmicity in DLPFC and NAc underlying the relationships between circadian rhythms, sleep, and OUD. Further, our results suggest that treatments to target biological pathways altered in subjects with OUD may be more effective when given at the time of day when the alteration is most robust. Thus, our insights will hopefully provide the opportunity for new therapeutics in the treatment of OUD.

## Acknowledgments

We would like to thank the staff and technicians who work diligently as part of the Brain Tissue Program at the University of Pittsburgh. Human tissue was obtained from the NIH NeuroBioBank and the University of Pittsburgh Brain Tissue Donation Program. This study was funded by the Hamilton Family Prize for Basic Neuroscience Research in Psychiatry at the University of Pittsburgh School of Medicine to R.W.L.; NIDA F30DA053020 to B.N.P.; NIDA DP1DA046585 to A.R.P.; NHLBI R01HL150432 to R.W.L.; and NIDA R01DA051390 to M.L.S. and R.W.L.

