## Supplementary Materials for "Molecular rhythm alterations in prefrontal cortex and nucleus accumbens associated with opioid use disorder"

Running Title: Transcriptional Rhythms in OUD

Xiangning Xue B.S.^1^, Wei Zong B.S.^1^, Jill R. Glausier Ph.D.^2^, Sam-Moon Kim Ph.D.^2,3^, Micah A. Shelton M.S.^2^, BaDoi N. Phan^4^, Chaitanya Srinivasan^4^, Andreas R. Pfenning Ph.D.^4,5^, George C. Tseng Sc.D.^1^, David A. Lewis MD^2^, **Marianne L. Seney Ph.D.^2,3^*,** and **Ryan W. Logan Ph.D.^6,7^***

^1^Department of Biostatistics, University of Pittsburgh, Pittsburgh, PA 15261, USA.

^2^Translational Neuroscience Program, Department of Psychiatry, University of Pittsburgh School of Medicine, Pittsburgh, PA 15219, USA.

^3^Center for Adolescent Reward, Rhythms, and Sleep, University of Pittsburgh, Pittsburgh, PA 15219, USA.

^4^Department of Computational Biology, Carnegie Mellon University, Pittsburgh, PA 15213, USA.

^5^Neuroscience Institute, Carnegie Mellon University, Pittsburgh, PA 15213, USA.

^6^Department of Pharmacology and Experimental Therapeutics, Boston University School of Medicine, Boston, MA 02118

^7^Center for Systems Neuroscience, Boston University, Boston, MA 02118

***To whom correspondence should be addressed:**

Ryan W. Logan, PhD

Department of Pharmacology and Experimental Therapeutics

Boston University School of Medicine

700 Albany Street

Boston, MA 02118

617-358-9563

Marianne L. Seney, PhD

Department of Psychiatry

University of Pittsburgh School of Medicine

450 Technology Drive

412-624-3072

**SUPPLEMENTARY TABLES**

**Table S1.** Subject summary demographic and tissue characteristics

| **Characteristic** | **Unaffected comparison (n=20)** | **Opioid**  **dependent**  **(n=20)** |
| --- | --- | --- |
| Age | 47.3 ± 9.5 | 46.9 ± 7.3 |
| Sex | 10M 10F | 10M 10F |
| Race | 13W 7B | 19W 1B |
| PMI (hours) | 15.7 ± 6.1 | 16.0 ± 5.3 |
| Brain pH | 6.6 ± 0.3 | 6.4 ± 0.2 |
| RIN | 8.0 ± 0.7 | 7.8 ± 0.7 |
| Tissue Storage Time (months) | 100.3 ± 86.1 | 103.0 ± 59.7 |
| TOD | 7.7 ± 7.3 | 8.6 ± 3.9 |

Values are mean ± SD. B, black; F, female; M, male; TOD, time of death; W, white

**SUPPLEMENTARY FIGURES**

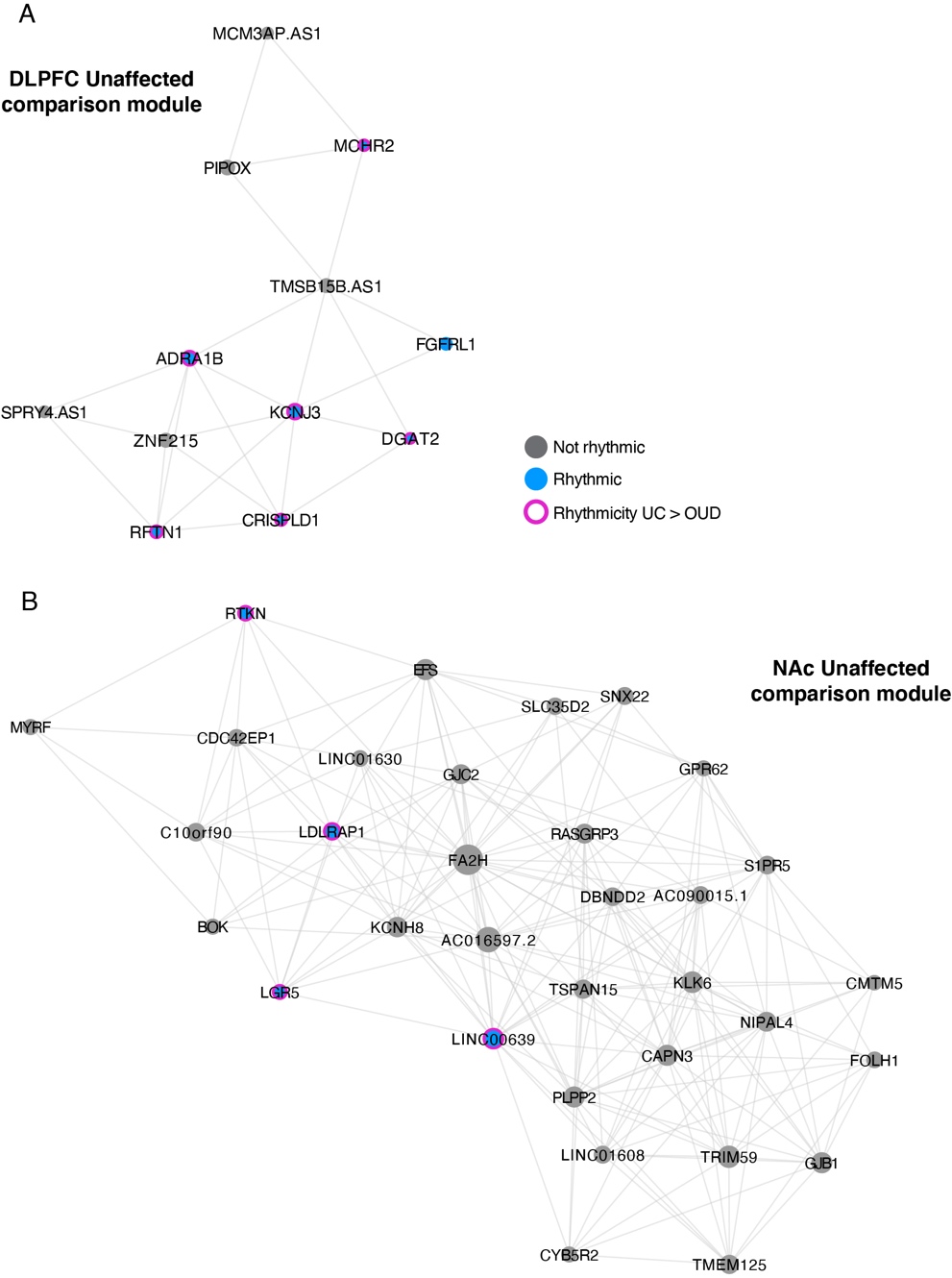

**Figure S1. Gene networks in the DLPFC and NAc of unaffected comparison subjects. A.** Weighted gene co-expression network analysis (WGCNA) was used to generate co-expression modules, with the network structure generated on each brain region separately. The identified modules that survived module preservation analysis were arbitrarily assigned colors and module differential connectivity (MDC) analysis compared the identified modules in OUD and unaffected comparison (UC) subjects. MDC analysis indicated a loss of connectivity in OUD subjects for one module in the DLPFC (**A**) and one module in the NAc (**B**). Node size indicates the degree of connectivity for that transcript. Blue nodes indicate rhythmic transcripts and magenta halos indicate transcripts that were more rhythmic in UC subjects compared to subjects with OUD. Edges indicate significant co-expression between two particular transcripts.
